# Overexpressing FLP Recombinase Can Cause Synaptic Trafficking Defects in *C. elegans*

**DOI:** 10.64898/2026.10.01.756008

**Authors:** Zephyr Lenninger, Frederic J. Hoerndli

## Abstract

Site-specific recombinases, such as Cre recombinase (Cre) and FLP recombinase (FLPase), are popular tools that enable precise spatiotemporal control over fluorescent reporter genes. However, studies show that increased Cre expression can lead to cytotoxic phenotypes including DNA damage, cell cycle arrest, cell death, and aneuploidy. In contrast, there is little-to-no known cytotoxicity associated with FLPase. FLPase is considered to be the safer, if less efficient, recombinase. To compensate for decreased efficiency, researchers will often increase expression of the FLPase to ensure a complete gene edit. Despite previous evidence that FLPase is nontoxic at high doses, we report that high expression of FLPase from multi-copy extrachromosomal arrays can disrupt synaptic protein trafficking in *C. elegans*. We discovered several off-target effects of high FLPase expression including mislocalization of the pre-synaptic marker FLPon::mScarlet::SYD-2 and decreased synaptic abundance, transport, and expression of the ionotropic glutamate receptor subunit GLR-1. Neither overexpression of an inactive FLPase, low activity FLPase, nor mCherry under the same promoter elicited these effects. However, single-copy expression of the FLPase construct reliably tags FLPon::mScarlet::SYD-2 and does not lead to these off-target effects. This work supports growing awareness that tyrosine recombinases can become cytotoxic at high doses.

## Introduction

Site-specific recombinases, such as Cre recombinase (Cre) and FLP recombinase (FLPase), recognize and bind specific target sequences (e.g., loxP, FRT) to catalyze DNA rearrangements (Broach and Hicks 1980; Sternberg and Hamilton 1981). In fluorescent microscopy, recombinase systems (Cre/lox; FLP/FRT) enable precise, spatiotemporal control over fluorescent reporter genes engineered to include these target sequences. Researchers can visualize endogenous proteins in specific tissues by strategically inserting target sequences into the genome while driving recombinase expression under tissue specific promoters (Livet et al. 2007; Jensen and Dymecki 2014). These enzymes perform without additional cofactors, leading to efficient genome modifications (Grindley et al. 2006). However, there is growing evidence that recombinase expression can become cytotoxic.

Cre is known to have harmful side effects in non-human primates (Yoshimatsu et al. 2023), mice (Higashi et al. 2009; Bersell et al. 2013; Bohin et al. 2018; Rezai Amin et al. 2019; Li et al. 2023; Straube et al. 2023; Liffers et al. 2025), Drosophila (Nern et al. 2011; Frickenhaus et al. 2015), enterocyte organoids (Moll et al. 2022), and mammalian cell lines (Loonstra et al. 2001; Patsch et al. 2010; Janbandhu et al. 2014). These studies report cytotoxic phenotypes including DNA damage, cell cycle arrest, cell death, and aneuploidy. In tissue, these effects manifest as cardiac dysfunction (Wang et al. 2020; Rashbrook et al. 2022; Li et al. 2023), gastrointestinal disruption (Jae Huh et al. 2010), and neurotoxicity (Forni et al. 2006; Nern et al. 2011; Rezai Amin et al. 2019; Erben et al. 2022). The cytotoxic effects of Cre occur at high doses and are thought to be due to DNA damage from aberrant endonuclease activity at pseudo-loxP sites (Rashbrook et al. 2022). Though FLPase and Cre are both tyrosine recombinases (Grindley et al. 2006), there is little-to-no known cytotoxicity associated with FLPase and its variants (FLPe, FLPo). In Drosophila, high expression of FLPase did not cause the same developmental toxicity seen with Cre (Nern et al. 2011; Frickenhaus et al. 2015). In cell culture FLPase expression did not show the cell proliferation defects, cell-cycle arrests, or increases in DNA repair machinery observed with Cre expression (Patsch et al. 2010; Galla et al. 2011). Indeed, FLPase is widely considered to be a safer, if less efficient, alternative to Cre. The ability to safely overexpress FLPase is valuable because increasing recombinase expression increases recombinase efficiency (Kondo et al. 2009; Patsch et al. 2010). Indeed, it is common to intentionally overexpress FLPase to increase the fluorescence of FLPon reporters. Existing methods for implementing FLPase optimize for thermostability and recombination fidelity but neglect to evaluate dose-dependent cellular phenotypes beyond broad defects in cell proliferation or viability. Addressing this gap is imperative to those using FLPon systems to tag endogenous proteins because minor levels of DNA damage can cause disruptions to protein trafficking without leading to cell death (Farber-Katz et al. 2014). No study has systematically examined whether high levels of FLPase can affect protein trafficking or homeostasis. This is particularly relevant given that multi-copy transgenic approaches are common methodologies across models for increasing FLPase efficiency and can yield enzyme concentrations orders of magnitude higher than single-copy systems. It remains unknown whether elevated FLPase expression disrupts proteostasis, membrane trafficking, or localization of synaptic proteins- processes that are highly sensitive to cellular stress. *C. elegans*, with its genetic tractability and transparent anatomy, provides an ideal platform to detect off-target effects of high FLPase expression *in vivo*, especially using neuronal synaptic proteins whose trafficking is sensitive to disruptions in neuronal homeostasis.

In this study, we directly compare the impact of multi-copy and single-copy FLPase transgenes on synaptic protein trafficking and localization in *C. elegans*. We implement a dual-genetic approach to mark pre- and post-synaptic compartments in AVA- a set of command interneurons involved in many well-defined behaviors such as navigation, avoidance (Maricq et al. 1995; Gray et al. 2005; Piggott et al. 2011) and olfactory associative learning (Hadziselimovic et al. 2014; Jin et al. 2016). Using this system, we uncover several off-target effects of overexpressed FLPase, including mislocalized synaptic markers and impaired GLR-1 transport. These phenotypes do not occur when FLPase is limited to a single copy, an inactive FLPase is overexpressed, or a less active FLPase is overexpressed, indicating that recombinase dose and activity are critical determinants for the off-target effects. Our study provides documentation that high FLPase expression can disrupt synaptic protein trafficking in *C. elegans* command interneurons. In addition, we also identify genomic single-copy insertions of FLPase as a practical mitigation strategy for the off-target effects.

## Results

The AVA command interneurons have defined regions of pre- and post-synaptic compartments that were identified and mapped by electron microscopy (EM) (Cook et al. 2019). To label pre-synaptic terminals in AVA, we utilize the FLP/FRT system with the D5 variant of FLPase (Nern et al. 2011; Schwartz and Jorgensen 2016) to tag endogenous SYD-2 with mScarlet (Figure 1a) (McDonald et al. 2023). To visualize glutamatergic post-synaptic compartments in AVA, we express a single-copy genetic replacement of GLR-1, the homologue of human GluA1/GluA2, tagged with super-ecliptic pHluorin (SEP) (Sankaranarayanan et al. 2000) to represent the population of GLR-1 at the cell surface (Figure 1b). We have shown that this single-copy SEP::GLR-1 is sufficient to rescue behavioral sensitization (Knight et al. 2026 June 12). By imaging animals in a ventral position, we can visualize AVA synaptic regions (Figure 1c). We assess the impact of FLPase expression in AVA by comparing two transgenic approaches that vary in copy number. For a high-dose FLPase, we use extrachromosomal arrays which can carry hundreds of copies of a transgene (Stinchcomb et al. 1985; Mello et al. 1991). Due to the variability of arrays, we use 3-5 independent transgenic lines in experiments where GLR-1 is quantified (Figure 3-4; S3). For a low-dose FLPase, we used rRMCE (Nonet 2023) to insert a single copy FLPase into the genome. We imaged the impact of multi-copy and single-copy FLPase on these synaptic markers and found that marker localization differed depending on the dose of FLPase (Figure 1d-e).

**Figure 1.**
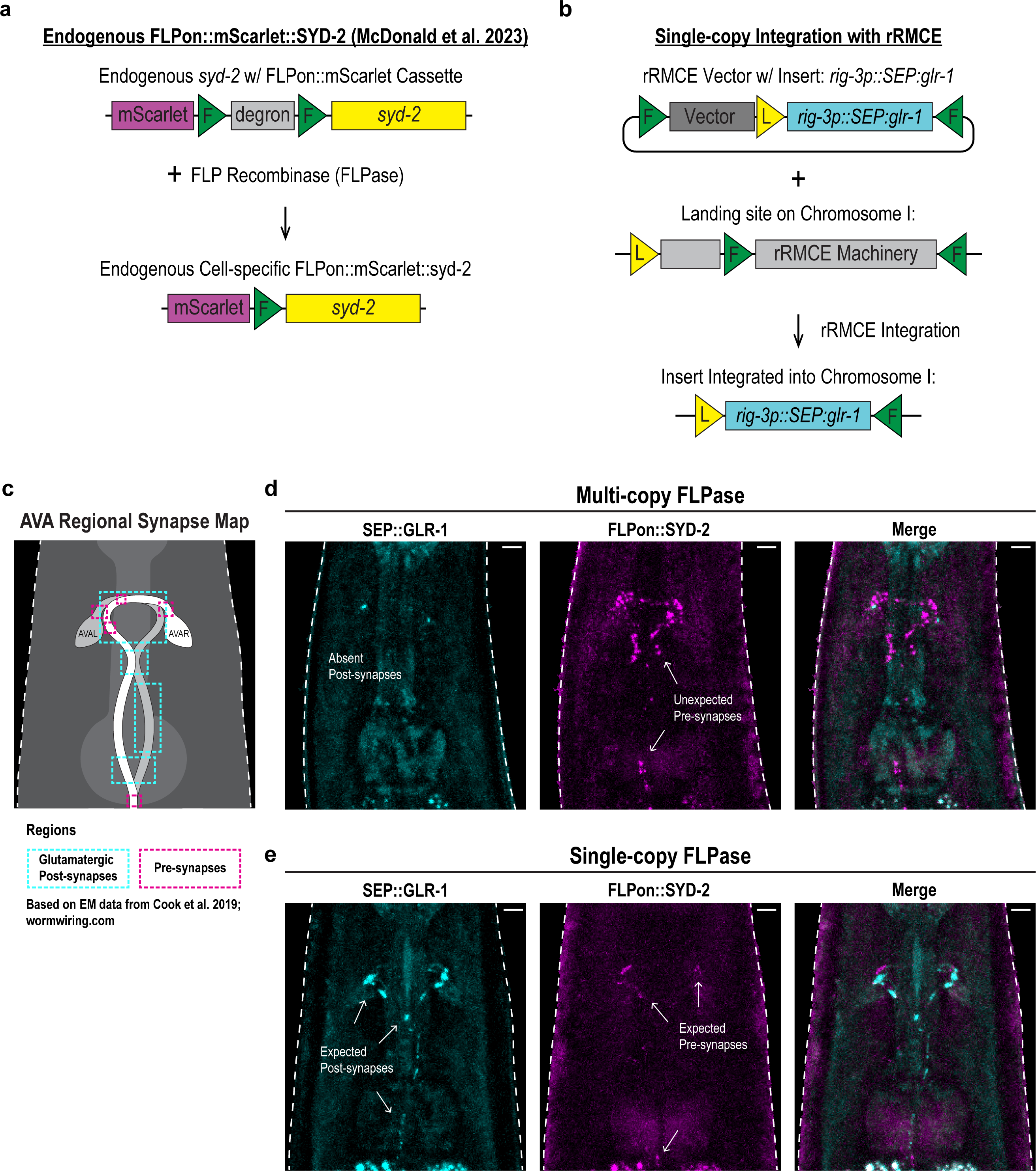
FLPase expression alters synaptic marker distribution. **a)** Diagram of FLPon::mScarlet::*syd-*2 (McDonald et al., 2023). **b)** Diagram of single-copy SEP::GLR-1. **c)** Illustration of AVA interneurons with a ventral view of synaptic regions in AVA; boxes denote regions containing pre- and/or post-synapses based on electron microscopy (Cook et al., 2019). **d, e)** Representative images at 40x of SEP::GLR-1 (cyan) and FLPon::mScarlet::SYD-2 (magenta) when **d)** FLPase is expressed from a multi-copy extrachromosomal array (FJH 972) and **e)** when FLPase is expressed from a single-copy genomic insertion (FJH 976). Scale bars: 5µm. FJH strain details are listed in Table 1.

Overexpressing FLPase in AVA causes FLPon::mScarlet::SYD-2 puncta to localize to regions that do not correspond to known pre-synaptic compartments in AVA (Figure 1c-d). To better analyze this, we captured a region of AVA called the bifurcation that has a distinct segregation of pre-synapses posterior to the point where the two AVA neurites join to enter the ventral nerve cord (Figure 2a). We took a line scan of the 80 pixels (∼8 um) directly anterior (upper branch) and posterior to the bifurcation (distance = 0 px), and we compared multi-copy and single-copy FLPase expression. Both the line scan and the images clearly show the presence of ectopic FLPon::mScarlet::SYD-2 puncta anterior to the bifurcation in multi-copy FLPase animals (Figure 2b-c). The expected pre-synaptic distribution is visibly represented in animals with single-copy FLPase (Figure 2d-e). Therefore, FLPase overexpression may misrepresent FLPon::mScarlet::SYD-2 localization in AVA. We attempted to image transport of FLPon::mScarlet::SYD-2 and quantify FLPon::mScarlet::SYD-2 in AVA cell bodies but could not detect either transport or somatic expression of SYD-2 in either multi- or single-copy FLPase strains (data not shown).

**Figure 2.**
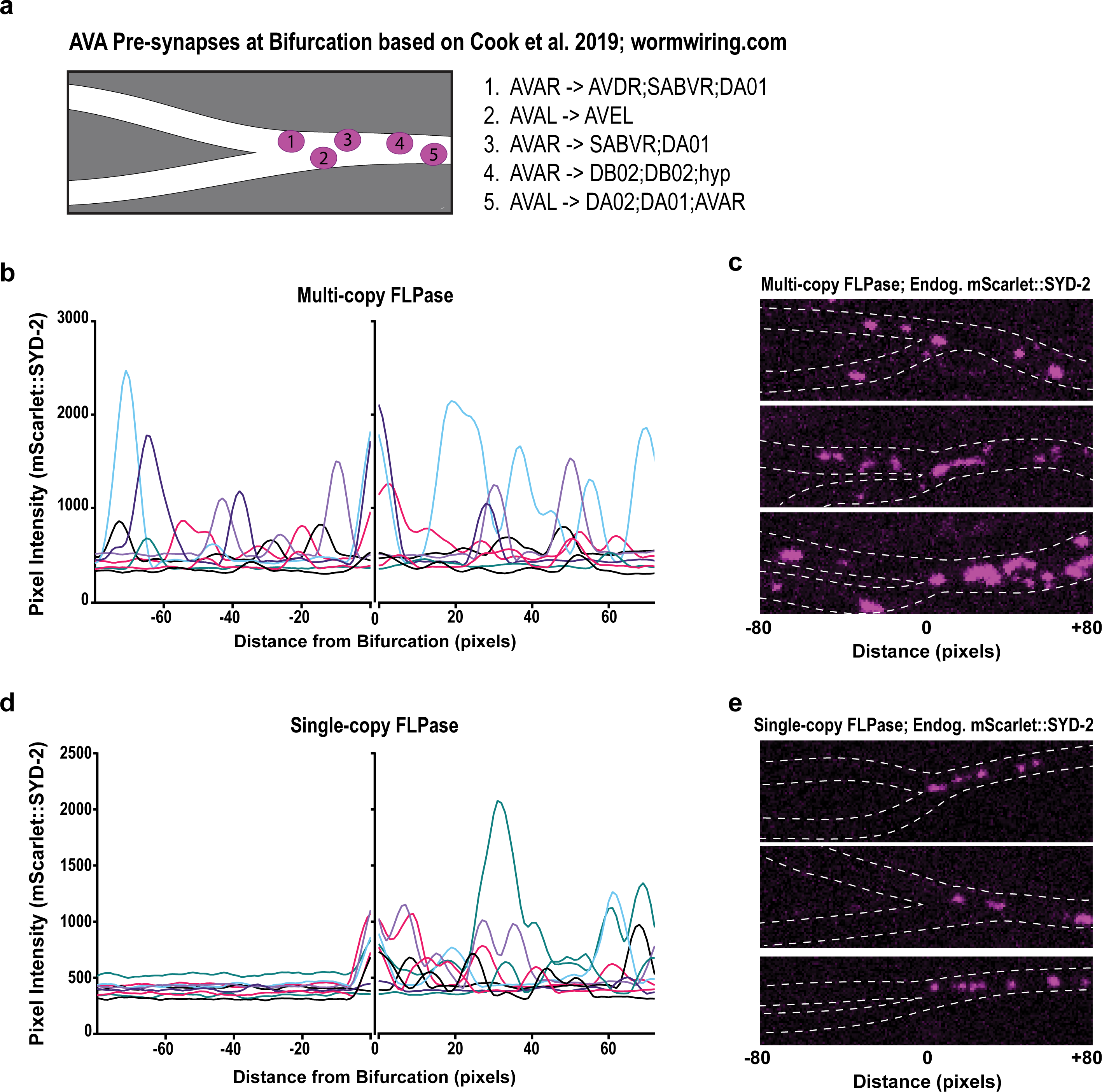
FLPase overexpression misrepresents endogenous pre-synaptic marker FLPon::mScarlet::SYD-2 in AVA. **a)** Illustration of pre-synapses at the bifurcation of AVA based on electron microscopy (Cook et al., 2019) with labels denoting [pre-synaptic neuron -> post-synaptic cell], not drawn to precise scale. **b)** FLPon::mScarlet::SYD-2 intensity from line scans oriented from the branching point of the bifurcation (0 px) in animals expressing high levels of FLPase from a multi-copy extrachromosomal array (n= 8 animals; FJH 972). **c)** Three replicate representative images of FLPon::mScarlet::SYD-2 puncta at bifurcation in AVA from animals with the FLPase multi-copy array. **d)** FLPon::mScarlet::SYD-2 intensity from line scans oriented from the branching point of the bifurcation (0 px) in animals expressing single-copy genomic FLPase (n= 9 animals; FJH 976). **e)** Three replicate representative images of FLPon::mScarlet::SYD-2 puncta at bifurcation in AVA from animals with single-copy FLPase. 80 pixels = ∼8 um. FJH strain details are listed in Table 1.

The absence of SEP::GLR-1 puncta when FLPase is overexpressed (Figure 1d) suggests a defect in GLR-1 surface localization which could reflect a broader trafficking deficit. A key step in GLR-1 trafficking is active transport by kinesin-1 from the cell-bodies to the synapses (Hoerndli et al. 2013). To determine if overexpressing FLPase to visualize FLPon::mScarlet::SYD-2 in AVA can impact long-distance GLR-1 transport, we overexpressed FLPase in a GLR-1::GFP imaging background. As previously described (Hoerndli et al. 2013; Doser et al. 2024), we recorded streams of GLR-1::GFP transport events *in vivo* and quantified intensity, number, and speed of transport events.

We found that overexpressing FLPase negatively impacts GLR-1 trafficking. Animals overexpressing FLPase have a significant decrease in the amount of GLR-1::GFP transported (Figure 3a-b), number of transport events (Figure 3c-e), and speed of those events (Figure 3f-g). These data suggest a pleiotropic effect on trafficking that could lead to somatic buildup of GLR-1. To address this, we quantified GLR-1::GFP in the AVA cell bodies. In animals overexpressing FLPase, we observe a decrease in GLR-1::GFP signal in the cell bodies (Figure S1a-b). This suggests that overexpressing FLPase likely impacts multiple steps of the GLR-1 trafficking pathway, including somatic expression of GLR-1.

**Figure 3.**
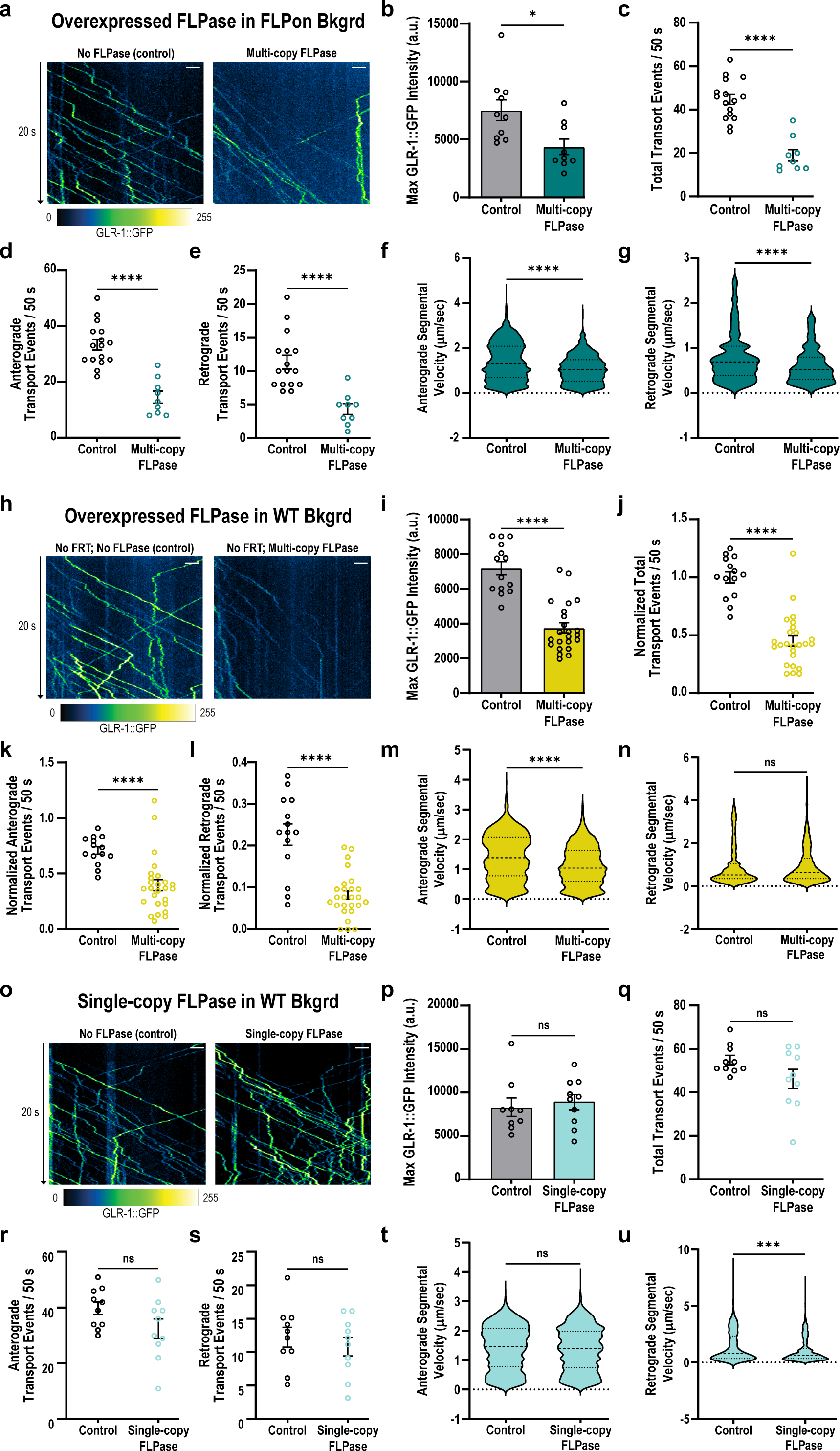
FLPase overexpression impairs GLR-1 trafficking in AVA. **a)** Kymographs showing GLR-1::GFP transport events in AVA when FLPase is not expressed (control) or is overexpressed (multi-copy FLPase). **b)** Quantified global maximum intensities of 50s long kymographs. **c-e)** Number of **c)** total, **d)** anterograde, and **e)** retrograde transport events from 50s long kymographs. **f-g)** Segmental velocity in **f)** anterograde and **g)** retrograde directions. n=9-16 animals each condition (FJH 817 and 877). **h)** Kymographs showing GLR- 1::GFP transport events in AVA when FLPase is not expressed (control) or is overexpressed (multi-copy FLPase) in a no-FRT background. **i)** Quantified global maximum intensities of 50s long kymographs. **j-l)** Number of **j)** total, **k)** anterograde, and **l)** retrograde transport events from 50s long kymographs. **f-g)** Segmental velocity in **f)** anterograde and **g)** retrograde directions. n= 14-26 animals for each condition (FJH 737, 991-963; five replicate strains, normalized to same day controls). **o)** Kymographs showing GLR-1::GFP transport events in AVA when FLPase is not expressed (control) or is expressed as a single copy. **p)** Quantified global maximum intensities of 50s long kymographs. **q-s)** Number of **q)** total, **r)** anterograde, and **s)** retrograde transport events from 50s long kymographs. **t-u)** Segmental velocity in **t)** anterograde and **u)** retrograde directions. n= 10 animals for each condition (FJH 737 and 934). For all kymographs, anterograde transport is represented by traces moving left to right, time is represented on the y-axis, and distance on the x-axis. Statistical significance was determined using a Mann-Whitney test. *p<0.05, **p<0.01, ***p<0.001, ****p<0.0001, ns: not significant. Error bars represent mean ± s.e.m. Scales bars are 2µm. For kymographs, LUT: greenfireblue. FJH strain details are listed in Table 1.

Since SYD-2 is a critical synaptic scaffold, it is possible that the trafficking defects are off-target effects of the FLPon mScarlet tag fused to SYD-2, which could disrupt its function. To test this possibility, we created a recombinase-only control by overexpressing FLPase without a FLPon background. Despite the absence of FRT sites or tagged SYD-2, we observe a significant decrease in quantity of transported GLR-1::GFP (Figure 3h-i) and number of transport events (Figure 3j-l) across five replicate strains. In addition, we observe a decrease in speed of anterograde, but not retrograde, transport events (Figure 3m-n). This suggests that high expression of FLPase is sufficient to induce GLR-1 transport defects in the absence of a FLPon cassette or FRT target sites and that these off-target effects persist across distinct strains with independently formed transgenic arrays.

Given that animals expressing single-copy FLPase displayed normal SEP::GLR-1 puncta distribution (Figure 1e), we sought to determine if the GLR-1::GFP trafficking deficits could be alleviated by expressing FLPase at this lower level. We measured GLR-1::GFP transport in animals expressing single-copy FLPase and found that this does not impact the quantity of transported GLR-1::GFP (Figure 3o-p) or number of transport events (Figure 3q-s). The speed of anterograde events is also unaffected, but the speed of retrograde events is decreased (Figure 3t- u). It is important to note that retrograde transport events are infrequent, and these observations could be due to stochastic effects of the videos obtained. In addition, single-copy FLPase expression does not decrease GLR-1::GFP levels in the cell-bodies (Figure S1c-d).

To better understand the impact of the trafficking defect, we quantified mean intensity and area of FLPon::mScarlet::SYD-2 and SEP::GLR-1 puncta in AVA. The fluorescence of FLPon::mScarlet::SYD-2 is dependent on the presence of the FLPase, so we could not incorporate a no-recombinase control. Previous studies report that increasing recombinase dose can increase fluorescent signal from a FLPon reporter (Kondo et al. 2009; Patsch et al. 2010). However, we do not observe a significant change in FLPon::mScarlet::SYD-2 puncta intensity between multi-copy and single-copy FLPase conditions (Figure 4d). We do observe a significant increase in puncta area with multi-copy FLPase (Figure 4e). The SEP::GLR-1 fluorescence does not require FLPase activity. Yet, compared to a no-recombinase control, we observe a significant decrease in SEP::GLR-1 intensity and area when FLPase is overexpressed across three replicate strains (Figure 4f-g). We do not see this effect with the single-copy FLPase (Figure 4f-g). It is possible that overexpressing any construct could cause the SEP::GLR-1 defect. However, overexpressing mCherry under the same promoter does not impact SEP::GLR-1 (Figure S2). Further, we have published several examples of *glr-1* reporters under the *rig-3* promoter co-expressed with multi-copy *flp-18p* constructs that do not cause GLR-1 trafficking defects (Hoerndli et al. 2013; Hoerndli et al. 2015; Hoerndli et al. 2022; Stetak et al. 2024).

**Figure 4.**
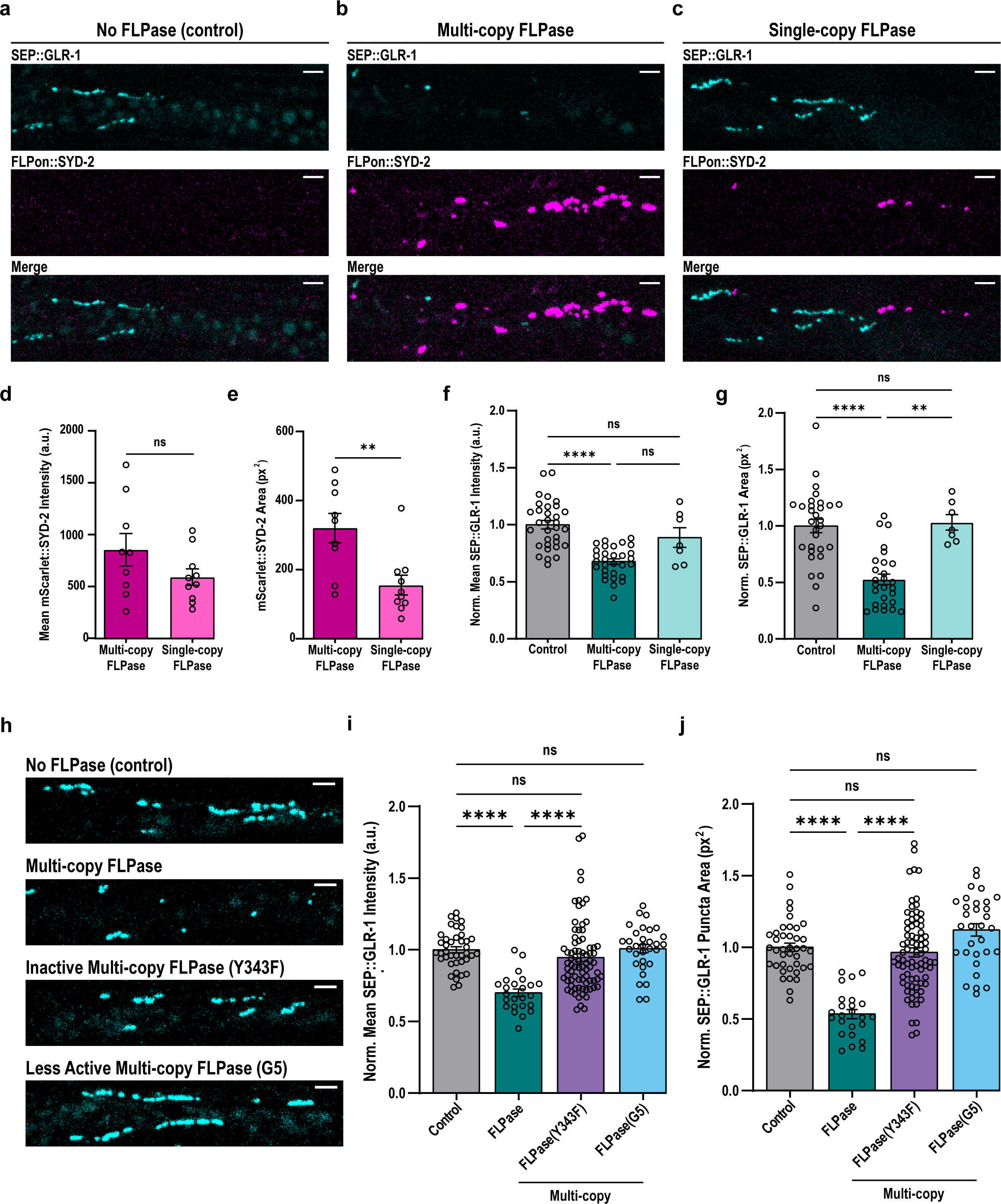
FLPase expression and activity impact synaptic SYD-2 and GLR-1 in AVA. a-c) Representative images of AVA bifurcation at 100x of SEP::GLR-1 (cyan) and FLPon::mScarlet::SYD-2 (magenta) when **a)** FLPase is not expressed (control), **b)** FLPase is overexpressed from a multi-copy extrachromosomal array, and **c)** FLPase is expressed from a single-copy genomic insertion. **d, e)** Quantification of FLPon::mScarlet::SYD-2 **d)** intensity and **e)** area. n=7-12 animals for condition (FJH 972 and 976); statistical significance determined with a Mann-Whitney test. **f, g)** Quantification of SEP::GLR-1 **f)** intensity and **g)** area. n=7-30 animals for each condition (FJH 880, 972, 960, 1140 and 976; three replicate strains normalized to same day controls); statistical significance determined with a Kruskal-Wallis’s test followed by Dunn’s multiple comparisons. **h)** Representative images of SEP::GLR-1 at AVA bifurcation when different FLPase variants are overexpressed. **i, j)** Quantification of SEP::GLR-1 **i)** intensity and **j)** area when FLPase, inactive FLPase(Y343F), and less active FLPase(G5) are expressed from multi-copy extrachromosomal arrays. n=23-78 animals for each condition (FJH 880, 972, 1078, 1073, 1074, 1079, 1075, 1141-1143; 3-5 replicate strains normalized to same day controls). Statistical significance determined with a Kruskal-Wallis test followed by Dunn’s multiple comparisons. **p<0.01, ****p<0.0001, ns: not significant. Error bars represent mean ± s.e.m. Scale bars: 2µm. FJH strain details are listed in Table 1.

It is possible that the SEP::GLR-1 deficit is a result of FLPase activity. To test this hypothesis, we mutated the catalytically active tyrosine (Y343F; Evans et al. 1990) in our FLPase construct. As expected, animals overexpressing FLPase(Y343F) did not exhibit FLPon::mScarlet::SYD-2 fluorescence. We did not see a decrease in SEP::GLR-1 mean or area across the five replicate strains (Figure 4h-j), though we did see a decrease in SEP::GLR-1 mean, but not area, in one out of the five isolates (Figure S3c-d). To further test the sensitivity of SEP::GLR-1 to FLPase activity, we utilized a less active FLPase variant (G5; Nern et al. 2011). Overexpressing FLPase(G5) did not impact SEP::GLR-1 mean nor area (Figure 5h-j) across three replicate strains (Figure S3e-f).

## Discussion

The absence of cytotoxic phenotypes is not proof of a reagent’s neutrality. Cre recombinase exhibits broad cytotoxic effects that translate to overt, observable defects corresponding to the cell-type in which the recombinase is expressed. In comparison to these broad cellular phenotypes, the deficits we found are subtle. The animals in this study show no obvious differences in development or lifespan, yet they exhibit significant defects in protein trafficking and localization. This subtlety is particularly insidious as it increases the risk that such effects could be misinterpreted as true biological phenomena. Our data suggest that high FLPase activity can disrupt protein trafficking in neurons by suppressing protein abundance, which indicates underlying cellular dysfunction from recombinase activity that has previously evaded detection.

Our work has parallels with a recent study of Cre toxicity in murine dopaminergic neurons that reports neural dysfunction and suppressed gene expression due to high doses of Cre (Erben et al. 2022). Given the similarity of phenotypes, it is possible that the model for Cre toxicity may describe a general mechanism of tyrosine recombinase toxicity that includes FLPase. This model suggests that Cre causes off-target DNA damage by binding to pseudo-loxp sites that naturally occur in the genome. Thousands of pseudo-loxP sites are documented in the mammalian genome (Semprini et al. 2007; Shultz et al. 2011), but there are no comprehensive studies with genome-wide searches for pseudo-FRT sites in *C. elegans*, just a single report of a pseudo-FRT site on chromosome V (Nonet 2024 Mar 3). We show that overexpressing FLPase can cause GLR-1 trafficking deficits in animals lacking engineered FRT sites, which suggests that pseudo-FRT sites may be more frequent in the *C. elegans* genome. Future studies should assess the prevalence of pseudo-FRT sites in the *C. elegans* genome to predict the impact of off-target FLPase activity.

To our knowledge, this study is the first to indicate that FLPase may share mechanisms associated with Cre toxicity in neurons. It will be pertinent for future studies to define the precise titration of FLPase expression that would ensure sufficient activity for a complete gene edit without risking toxicity. We suspect that the dose will need to be tailored to each context, as differences in cell type and promoter strength will vary the outcome. Our data are limited in that they are restricted to the impact on *syd-2* and *glr-1* transgenes in AVA. Nevertheless, we provide proof-of-concept that FLPase expression can disrupt protein trafficking. We recommend the following strategies to mitigate this concern: 1. Trial transient, low expression, and low activity recombinase conditions. 2. Validate localization of FLPon markers with an independent reference or a split fluorophore approach. 3. Incorporate no-recombinase and recombinase-only controls whenever possible.

## Methods

*C. elegans* strains were maintained under standard conditions at 20 °C on NGM plates seeded with OP50. A detailed description of strains used in this study listed by figure is provided in Table 1. Transgenic animals were created following standard methods to form extrachromosomal arrays via microinjection (Mello et al. 1991) or single-copy insertions via rRMCE (Nonet 2023). For multi-copy extrachromosomal arrays, FLPase plasmids (pZL24; pZL28; pZL34) were injected at 20 ng/ul with the co-injection marker pCT61 (20 ng/ul) and pBSKs (40 ng/ul). All plasmids were created using In-Fusion Cloning (Takara Bio) unless otherwise specified. The details of each plasmid are described in Table 1. All constructs were verified via full plasmid sequencing through Plasmidsaurus (plasmidsaurus.com).

Imaging was performed using a Yokogawa CSUX1 spinning disc on an Olympus IX83 confocal microscope with 488 and 561 nm diode lasers (OPSL 50mW each; Andor ILE Laser Combiner). Images were acquired with Plan S-APO x40/0.95NA air or a x100/1.40NA oil objectives (Olympus) as specified and an Andor iXon Ultra EMCCD (DU-867) camera. All devices were controlled remotely for image acquisition using MetaMorph 7.10.1 (Molecular Devices). All animals were imaged as one-day-old adult hermaphrodites. Individual animals were placed in 1.6 µL of a 1:1 solution of either 30 mM muscimol (Cat# 195336, MP Biomedicals) or 100 mM sodium azide (Cat # 190380050, Acros Organics) in M9 to polystyrene beads (Polybead, Cat# 00876-15, Polysciences Inc) on a 10% agarose (Cat # 97064-134, VWR Chemicals) pad. A coverslip was placed on top of the pad, restraining the animal. The coverslip was manually adjusted to orient the ventral side of the animal to face the objective for optimal visualization of AVA (Hoerndli et al. 2013; Doser et al. 2023). The specific imaging parameters for each figure are described in Table 1.

To image GLR-1 transport, AVA neurites were located using the 488 nm excitation laser and x100 objective. A Z drift compensator (Olympus, IX3-ZDC2) was used to hold continuous focus. A proximal portion of the neurites was photobleached using a 2 W, 488 nm Coherent solid-state laser (Genesis MX MTM; 0.5 W output; 1 s pulse). The photobleached region was defined in MetaMorph using a Mosaic II digital mirror device (Andor Mosaic 3). An image stream was collected directly after photobleaching. Kymographs were generated as described in Hoerndli et al. 2013 using MetaMorph’s Kymograph tool. Transport events were counted manually from the kymographs. To quantify the change in fluorescence intensity of the transport events, we plotted the maximum intensity of the whole kymograph because the mean would be skewed by the significant decrease in number of transport events. Segmental velocity was quantified using the ImageJ plugin KymoAnalyzer (Neumann et al. 2017).

AVA cell bodies were located using the 488 nm excitation laser and x100 objective. Image stacks were converted to maximum intensity projections using FIJI (ImageJ). ROIs were manually drawn around the cell bodies to quantify mean fluorescence intensity.

Synapses at the AVA bifurcation were located using the 488 nm excitation laser and x100 objective. Image stacks were converted to maximum intensity projections using FIJI (ImageJ). For SYD-2 distribution (Figure 2), line scan intensity profiles were acquired in FIJI using a 1- pixel-wide segmented line. Line scans originated at the bifurcation branching point and extended independently in the anterior and posterior directions from the common origin point. Intensity data were smoothed using a second-order polynomial function across a 5-neighbor window to reduce noise while preserving puncta peak structure. For quantification of mean fluorescence intensity and area, regions of interest (ROIs) were drawn manually around the neurites based on morphology and measurements were limited to threshold (MaxEntropy algorithm) within each ROI to isolate puncta from background without bias.

Representative images were chosen to represent the mean. All images were processed using FIJI (ImageJ). For presentation, background subtraction was performed using FIJI’s rolling-ball algorithm with a radius of 40 px at each z-plane before max-projection. Linear adjustments of brightness and contrast were set based on the control and applied equally across compared conditions. No non-linear adjustments were made. Graphs were generated using GraphPad Prism. Multiple independent transgenic lines for an overexpressed construct were examined to confirm the effect of the extrachromosomal arrays. In these cases, the data were normalized to same day controls. Individual isolates are plotted in Figure S3. All data are represented as the mean ± the standard error of the mean. Illustrations of synaptic locations in AVA (Figure 1, Figure 2) are intended to be representative and are not drawn to scale; they are based on the N2U dataset from Cook et al. 2019 (wormwiring.org/apps/neuronMaps). Figures and illustrations were constructed in Adobe Illustrator (2024). Prism 10 (GraphPad) was used for statistical analysis and to generate graphs. Outliers were removed from datasets using the ROUT method (Q = 1%). Details on the number of animals and specific statistical tests are presented in the figure legends.

## Data availability

All plasmids and strains are available on request. All strains, plasmids, and reagents used in this manuscript are described in Table 1. Raw data is provided in Supplementary Table 1. All strains, reagents, and methods were used in accordance to our Institutional Biosafety Committee review under no 18-043B.

## Acknowledgements

We would like to thank the Caenorhabditis Genetics Center (CGC) supported by the National Institutes of Health - Office of Research Infrastructure Programs (P40 OD010440). We would also like to thank Michael Nonet for the kind gift of strains and plasmids as well as tips for the efficient implementation of rRMCE. We would also like to thank Nathan McDonald for gifting us the FLPon SYD-2 strains.

## Figure Legends

**Figure S1. Somatic GLR-1::GFP is reduced in AVA when FLPase is overexpressed.**

**a)** Representative images depicting intensity of GLR-1::GFP in the AVA cell bodies when FLPase is not expressed (control) or is overexpressed (multi-copy FLPase). **b)** Quantification of GLR-1::GFP intensity in the cell bodies of AVA when FLPase is overexpressed. n= 19-25 cells each condition (FJH 817 and 877). **c)** Representative images depicting intensity of GLR-1::GFP in the AVA cell bodies when FLPase is not expressed (control) or is expressed as a single copy. **f)** Quantification of GLR-1::GFP intensity in the cell bodies of AVA. n= 21 cells each condition (FJH 737 and 934). Statistical significance was determined using a Mann-Whitney test; ***p<0.001, ns: not significant. Error bars represent mean ± s.e.m. Scale bars: 2µm. For images, LUT: greenfireblue. FJH strain details are listed in Table 1.

**Figure S2. Overexpressing mCherry in AVA does not decrease GLR-1 abundance at synapses.**

**a, b)** Representative images of AVA bifurcation when mCherry is **a)** not expressed (control) or **b)** overexpressed from an extrachromosomal array. **c,d)** of SEP::GLR-1 **c)** intensity and **d)** area when mCherry is expressed from a multi-copy extrachromosomal array. n=18-21 animals for each condition (FJH 885 and 1081). Statistical significance was determined using a Mann-Whitney test; ns: not significant. Scale bars: 2µm. FJH strain details are listed in Table 1.

**Figure S3. Impact of individual isolates of FLPase extrachromosomal array on GLR-1 abundance at synapses.**

**a, b)** Quantification of SEP::GLR-1 **a)** area and **b)** intensity when FLPase is expressed from a multi-copy extrachromosomal array in three replicate strains n=7-30 animals each condition (FJH 880, 972, 960 and 1140). **c, d)** Quantification of SEP::GLR-1 **c)** area and **d)** intensity when FLPase(Y343F) is expressed from a multi-copy extrachromosomal array in five replicate strains. n=15-24 animals for each condition (FJH 880, 972, 1078, 1073, 1074, 1079, and 1075). **e, f)** Quantification of SEP::GLR-1 **e)** area and **f)** intensity when FLPase(G5) is expressed from a multi-copy extrachromosomal array in three replicate strains. n=8-17 animals for each condition (FJH 880, 1141-1143). Statistical significance from control was determined with a Kruskal- Wallis test followed by Dunn’s multiple comparisons. *p<0.05, **p<0.01, ***p<0.001, ****p<0.0001; unmarked comparisons to control are not significant. FJH strain details are listed in Table 1.

---

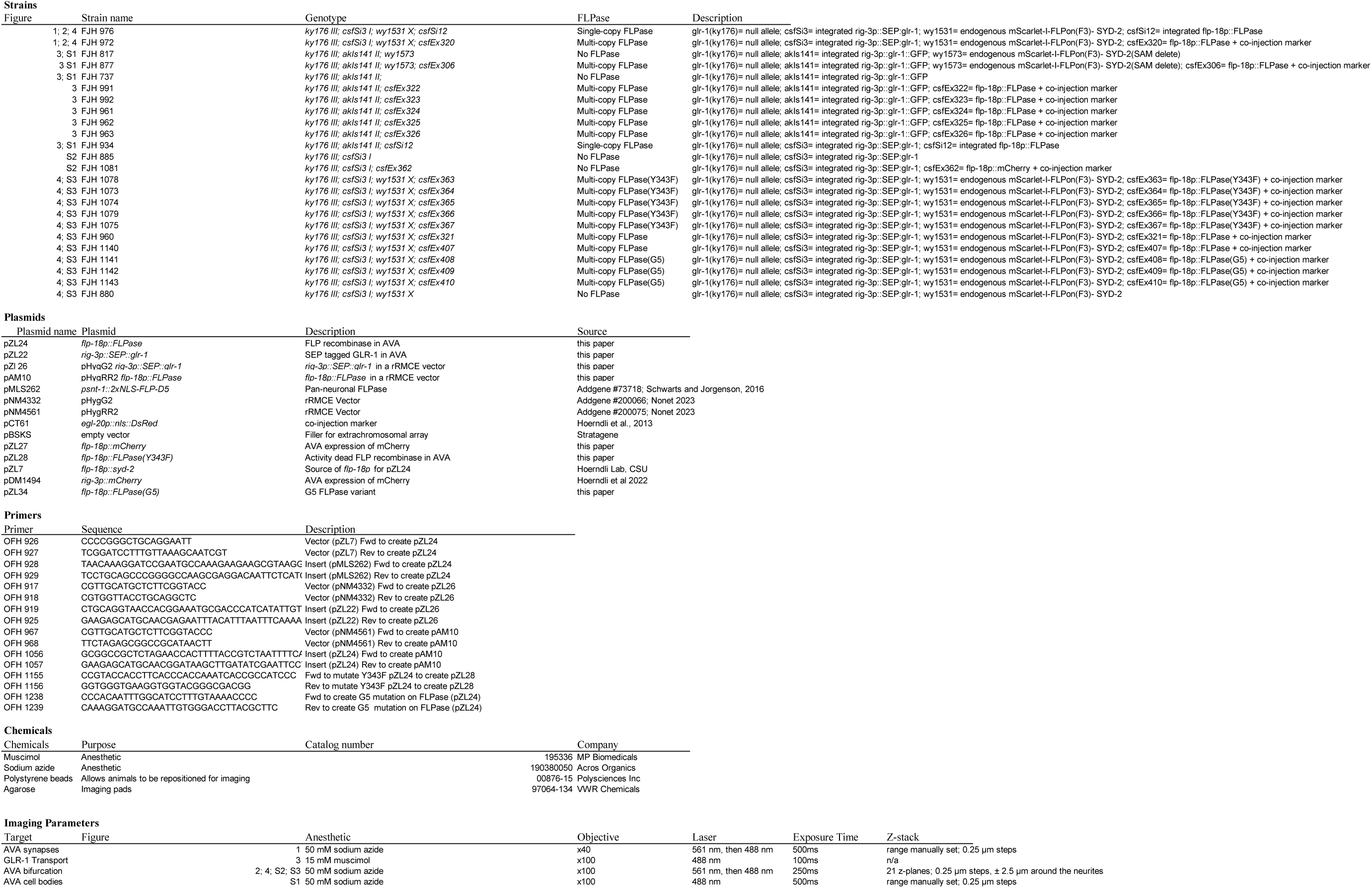

